# Effects of culture conditions on morphology and growth of *Linderina macrospora* (*Kickxellomycotina*, *Kickxellales*)

**DOI:** 10.64898/2026.07.30.741699

**Authors:** Tomohiko Ri, Teruhisa Masaki, Yousuke Degawa

## Abstract

The obscure life histories of many kickxellalean genera remain a bottleneck for comprehensive taxon sampling and phylogenetic reconstruction of the order. While *Kickxellales* has long been regarded as saprobes inhabiting soil or dung, the discovery of “amphibious fungi” such as *Unguispora*, which exhibits dimorphic growth between the animal gut and feces, suggests a cryptic gut-dwelling stage within these genera. Given its phylogenetic proximity to *Unguispora*, ecophysiological traits of *Linderina* were investigated to evaluate its potential association with the animal gut. Two isolates of *L. macrospora* were obtained from Japanese soil, representing the first record of this species in Japan. Physiological assays revealed that the optimal temperature for both vegetative growth and sporulation was 25–30 °C. Furthermore, comparative growth assays on different media demonstrated that sporocladium abundance per sporangiophore is sensitive to nutrient availability, and nutrient-poor media were determined to be the most suitable for evaluating morphological characterization. Under anaerobic, nutrient-rich conditions which are known to induce yeast-like growth in *Unguispora*, sporangiospores of *L. macrospora* produce arthrospores. Although marked morphological plasticity was observed during the arthrospore formation, the occurrence of yeast-like unicellular proliferation suggests a potential relationship with the animal gut. Additionally, vegetative growth and sporulation were markedly inhibited by white light exposure; notably, a lethal effect on growth was observed during incubation at 20 °C, indicating that the natural niche of the species is restricted to light-shielded environments. Our findings will help to elucidate the cryptic life cycles and evolutionary trajectories within *Kickxellales*.

## 1. Introduction

Recent molecular phylogenetic studies have led to a significant revision of the order *Kickxellales*, currently recognized as comprising one family and nine genera. Following the phylogenetic distinctions identified by Tretter et al. (2014), several taxa were reassigned: *Mycoëmilia* and *Spiromyces* within *Spiromycetales*, and *Ramicandelaber* within *Ramicandelaberales* (Doweld, 2014a, 2014b). Furthermore, the genus *Spirodactylon* was recently synonymized with *Coemansia* based on a nine-gene phylogenetic analysis (Reynolds et al., 2023). Despite these taxonomic advancements, a profound lack of fundamental biological insights remains the primary bottleneck, hindering the isolation of key lineages and the expansion of culture collections required for high-resolution phylogenetic inferences. Historically, kickxellalean fungi have been isolated from soil, dung, and other fungi; however, they are rarely encountered, except for *Coemansia* (Benny et al., 2016). For instance, *Dipsacomyces*, *Martensiomyces*, and *Pinnaticoemansia* have not been rediscovered since their original descriptions (Benjamin, 1961, 1959; Kurihara and Degawa, 2006). *Coemansia aureum* (*= Spirodactylon aureum*) and *Martensella* have only been reported from limited regions (Benny et al., 2001; Jackson and Dearden, 1948), and *Kickxella* is seldom found on vertebrate dung (Benny et al., 2014). Although Takashima et al. (2022) demonstrated that *Myconymphaea* can be stably isolated from the feces of *Lithobiomorpha* centipedes, the primary isolation sources and life histories of many other genera remain obscure.

The long-standing enigma surrounding the life histories of *Kickxellales* was fundamentally overturned by the discovery of *Unguispora*. Isolated from the excrement of camel crickets and crickets, this genus exhibits a dual life cycle comprising a saprobic stage on their feces and a symbiotic stage within their guts (Ri et al., 2022; Ri and Degawa, 2025). This finding suggests that other kickxellalean fungi, previously regarded strictly as soil or dung inhabitants, may similarly possess a life stage associated with the intestinal environment. Taxonomically, *Kickxellales* belongs to the subphylum *Kickxellomycotina*, which encompasses distinct saprobic, gut-dwelling, and mycoparasitic lineages (Benny et al., 2016). Prior to *Unguispora*, no fungus in this subphylum was known to bridge both saprobes and gut symbionts. Therefore, *Unguispora* represents a specialized ecological group termed “amphibious fungi,” which serves as an ecological model for illustrating the transition between saprobic and gut-dwelling lifestyles within *Kickxellomycotina* (Ri et al., 2022). Consequently, a comprehensive re-examination of the life histories of other genera is imperative to elucidate the evolutionary transitions within this subphylum.

*Unguispora* is known to be the sister to the genus *Linderina* (Ri and Degawa, 2025), which is also rarely isolated from soil and is characterized by dome-shaped, aseptate sporocladia (Raper and Fennell, 1952). *Linderina* currently comprises two species: *L. pennispora* and *L. macrospora*. *Linderina pennispora* produces many sporocladia on a sporangiophore and small sporangiola (17–21 × 3.5–4.5 μm) (Raper and Fennell, 1952), whereas *L. macrospora* produces one, or rarely two to three sporocladia on a sporangiophore and large sporangiola (28–38 × 3–5 μm) (Chang, 1967). *Linderina pennispora* is primarily documented in tropical regions, including Liberia, India, Malaysia, Taiwan, and Indonesia (Baijal, 1963; Ho et al., 2007; Kurihara et al., 2008; Loh et al., 2001; Raper and Fennell, 1952). On the other hand, *L. macrospora* exhibits a broader latitudinal range spanning Hong Kong, southeastern United States, Indonesia, and Taiwan (Chang, 1967; Chien, 1971; Chuang and Ho, 2009; Kurihara et al., 2008), which suggests its potential occurrence in Japan.

Given their phylogenetic proximity, we hypothesized that *Linderina* species are also amphibious fungi. One of the key indicators of this lifestyle is the ability to exhibit dimorphic growth. *Unguispora* typically produces hyphae and sporangiola in the saprobic phase on feces but switches to yeast-like growth within the host gut; this transition can be induced in vitro by incubating the sporangiola under anaerobic, nutrient-rich conditions that mimic the gut environment (Ri et al., 2022). In this study, we obtained two *Linderina* isolates from soil in Japan. These isolates were identified based on morphological characteristics and phylogenetic analyses, and their ability to undergo yeast-like growth was tested under various atmospheric and culture conditions. Furthermore, we investigated the effects of temperature, medium, and illumination on vegetative growth and sporulation to gain deeper insights into the distribution and life history of *Linderina*.

## 2. Materials and Methods

### 2.1. Isolation and morphological observation

Two isolates of *Linderina* were obtained from Japan. Of the two isolates, one was obtained from soil collected in a weedy area near a parking lot in Kuji-gun, Ibaraki prefecture on 16 Oct 2015. A 1:500 dilution of the soil sample was plated onto potato dextrose agar (PDA; Shimadzu Diagnostics Corp., Japan) supplemented with 100 μg mL^-1^ chloramphenicol (Fujifilm Wako Pure Chemical Corp., Japan) in the final volume. The plates were incubated at room temperature (ca. 25 °C) under indoor light. The other was isolated from soil collected in a mixed forest of needleleaf and broadleaf trees in Sakuragawa city, Ibaraki prefecture on 16 Apr 2021. The soil sample was incubated by the moist chamber method at room temperature under indoor light. Pure cultures were obtained by transferring their spore mass to PDA or half-strength malt-extract yeast-extract agar (1/2 ME-YE agar; Kurihara et al. 2000) with a fine needle and deposited to the National Biological Resource Center (NBRC) as NBRC 116291 and 116292. The isolates that had been incubated on 1/2 ME-YE agar or corn meal agar (CMA; Shimadzu Diagnostics Corp.) for 4–14d were observed for species identification and photographed using a SZ61 stereo microscope (Olympus, Japan) with an EOS kiss X10 (Canon, Japan) and a BX51 light microscope (Olympus) with differential interference with a DP73 digital camera (Olympus). Slides were mounted in lactic acid or lacto-fuchsin (Carmichael, 1955).

### 2.2 PCR, Sequencing

Fungal DNA was extracted from 2 wk old mycelia grown on 1/2 ME-YE agar according to the procedures described in Ri and Degawa (2025). Three nuclear loci were amplified: the 18S rRNA gene (SSU) using the primers SR1R/NS8Z (O’Donnell et al., 1998; Vilgalys and Hester, 1990), the internal transcribed spacer region (ITS) and 28S rRNA gene (LSU) using ITS1F/LR5 (Gardes and Bruns, 1993; Vilgalys and Hester, 1990). PCR amplification and sequencing were performed following Ri and Degawa (2025), and the resulting sequences were deposited in GenBank.

### 2.3. phylogenetic analyses

To elucidate the phylogenetic position of our isolates, sequences of the *Kickxellales* and *Orphellales* (as an outgroup) were sampled from the NCBI GenBank database based on Chuang et al. (2017) and integrated with our newly obtained sequences (Supplementary Table 1). Sequences for each locus were aligned using MAFFT v7.511 with default parameters (Katoh et al., 2019). Multiple sequence alignments were trimmed using the gappyout model in trimAl v1.2 (Capella-Gutiérrez et al., 2009). The trimmed alignments of the three loci were then concatenated for subsequent phylogenetic analyses. Maximum likelihood (ML) analysis was performed using IQ-TREE v3.0.1, with branch support assessed via 1000 ultrafast bootstrap (UFB) replicates (Hoang et al., 2018; Wong et al., 2026). The optimal nucleotide substitution model for each locus in the ML analysis was selected by ModelFinder (Kalyaanamoorthy et al., 2017) based on the corrected Akaike Information Criterion (AICc, Sugiura, 1978) and applied to a partitioned analysis (Chernomor et al., 2016): GTR+F+I+R3 for SSU, TIM2+F+I+G4 for ITS, and TIM3+F+I+G4 for LSU. Bayesian inference (BI) was conducted using MrBayes v3.2.7a (Ronquist et al., 2012). Optimal models for BI were determined based on the Bayesian Information Criterion (BIC, Schwarz, 1978) using Kakusan4 (Tanabe, 2011). The GTR+G model was selected for all three loci under a proportional model. Two independent, simultaneous Metropolis-coupled Markov chain Monte Carlo (MCMC) runs with four chains per run were executed for two million generations, sampling trees every 1000 generations. The average standard deviation of split frequencies (ASDSF) were calculated every 5000 generations. Convergence of the MCMC process was evaluated by ensuring an ASDSF < 0.01 and effective sample size (ESS) scores > 100 using MrBayes and Tracer v1.6 (Rambaut et al., 2018), respectively. The first 25% of the sampled trees were discarded as burn-in, and the remaining trees were used to construct a 50% majority-rule tree and determine posterior probabilities (PPs) for individual branches. All alignment datasets and phylogenetic trees are available in the Zenodo online repository (10.5281/zenodo.20374431).

### 2.4. Evaluation of the effects of cultural conditions on morphology and growth

To confirm yeast-like growth in *Linderina*, sporangiola of the isolates were incubated under the conditions that induce yeast-like growth in *Unguispora*, which are anaerobic conditions using AnaeroPouch-Anaero (Mitsubishi Gas Chemical Co., Inc., Japan) on 1/2 ME-YE agar at 20 °C (Ri et al., 2022). Each isolate was cultured in triplicate and monitored on a daily basis. For comparison, experiments were carried out with modified media using four types of media where each component excluding agar was omitted from 1/2 ME-YE agar (-ME; malt extract, -YE; yeast extract, -P; peptone, -G; glucose), PDA, and CMA. These experiments, including the conditions of 1/2 ME-YE agar, were also conducted at 25 °C. Additional cultures were incubated under microaerophilic conditions using AnaeroPouch-MicroAero (Mitsubishi Gas Chemical Co., Inc.) on 1/2 ME-YE agar at 20 °C.

Then, the isolates were incubated on CMA at 10, 15, 20, 25, 30, and 35 °C for 10 days in the dark by shading with aluminum foil to determine the optimal temperature for growth and sporulation. Each isolate was cultured in triplicate, and the colony size was determined by averaging the mean diameters (4 measurements per colony) of three replicates. The degree of sporulation was observed under the stereo microscope. Furthermore, the effects of medium and illumination conditions on growth and sporulation were evaluated. The media were classified into nutrient-rich (1/2ME-YE agar and PDA) and nutrient-poor (CMA and _LC_A (Miura and Kudo, 1970)) types. The illumination conditions were defined as either light or dark. The “light” treatments were exposed to continuous white LED light (30–40μmol m^-2^ s^−1^ photosynthetic photon flux density). The “dark” treatments were completely shaded using aluminum foil. The association between the effects of these conditions and temperature was also assessed varying at 20 and 25 °C. The experiment under 16 different conditions combining the above factors was conducted using triplicates of each isolate. Following a 10-day incubation period, colony diameters and colors, presence of sporulation, and number of sporocladia per sporangiophore were recorded. Simultaneously, the growth of vegetative hyphae and abundance of sporangiophores were visually graded using stereomicroscopy. A separate experiment was performed for NBRC 116291, where a white light condition with a 14-hour photoperiod was used for the treatments involving 1/2ME-YE agar and CMA media at 20 and 25 °C.

## 3. Results

### 3.1. Taxonomy

The morphology of our isolates is illustrated in Figure 1. Sporangiolum size, one of the key diagnostic characters, was 26–30.5(–32.5) × 3.5–5 μm, which fell within the range reported in the original description of *L. macrospora* (Chang, 1967). Other morphological features were broadly concordant with the type description, with some differences noted. These isolates produced wider pseudophialides and shorter sporangiola and sporangiospores (Fig. 1E, G, H) than the type description, where the sizes of pseudophialides, sporangiola, and sporangiospores are 10–17 × 2–3 μm, 26–38 × 3–5 μm, and 26–36 × 3–5 μm, respectively (Chang, 1967). Although the number of sporocladia per sporangiophore in our isolates varied with medium and isolate, it was more or less identical to that in *L. macrospora* isolates from Hong Kong, United States, and Taiwan (Chang, 1967; Chien, 1971; Chuang and Ho, 2009). Helicoid structures with thick walled, bright ochre-yellow cells in Indonesian isolates reported by Kurihara et al. (2008) were not observed in this study.

**Figure 1.**
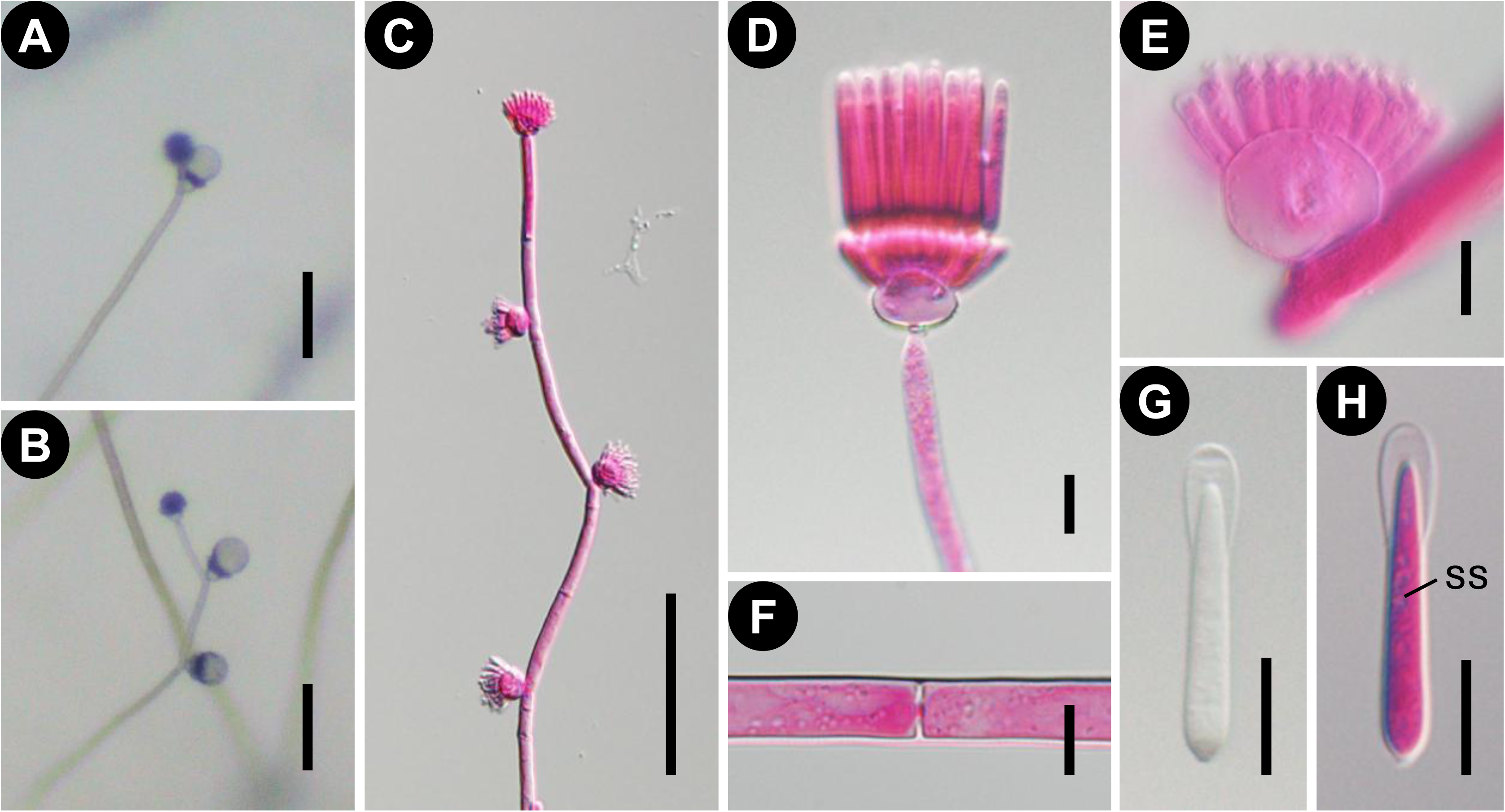
*Linderina macrospora* (NBRC 116291). **A, B** Stereo microscope. **C–H** Light microscope. **C–F, H** are mounted in lactic acid. **G** is mounted in lacto-fuchsin. **A–C** Apical portion of sporangiophores with sporocladia formed sympodially and acropetally. In mature, a droplet of sporangiola is formed on a sporocladium. **D** Sporangiola on pseudophialides born on the sporocladium. **E** Pseudophialides after detachment of sporangiola. **F** A septum of the sporangiophore with a central pore and a plug. **G, H** A sporangiospore (ss) within a sporangiolum. Scalebars: 100 μm (**A–C**); 10 μm (**D–H**).

ML and Bayesian phylogenetic analyses were performed using a combined dataset of 3323 nucleotides from three regions: SSU (1569 bp), ITS (803 bp), and LSU (951 bp). Of the aligned positions, 1489 sites were constant and 1834 were variable, including 1628 parsimony-informative sites. The Bayesian consensus tree was topologically congruent with the ML tree shown in Figure 2. Our isolates were grouped with other isolates of *L. macrospora*, including the ex-holotype strain, with robust support (100% ML UFB / 1.00 Bayesian PP; Fig. 2). Although the Indonesian isolates (ID05-F0180, ID05-F0214) formed a distinct lineage within the species, the deeper node encompassed all *L. macrospora* isolates with relatively low statistical support (70% ML UFB / 0.62 Bayesian PP).

**Figure 2.**
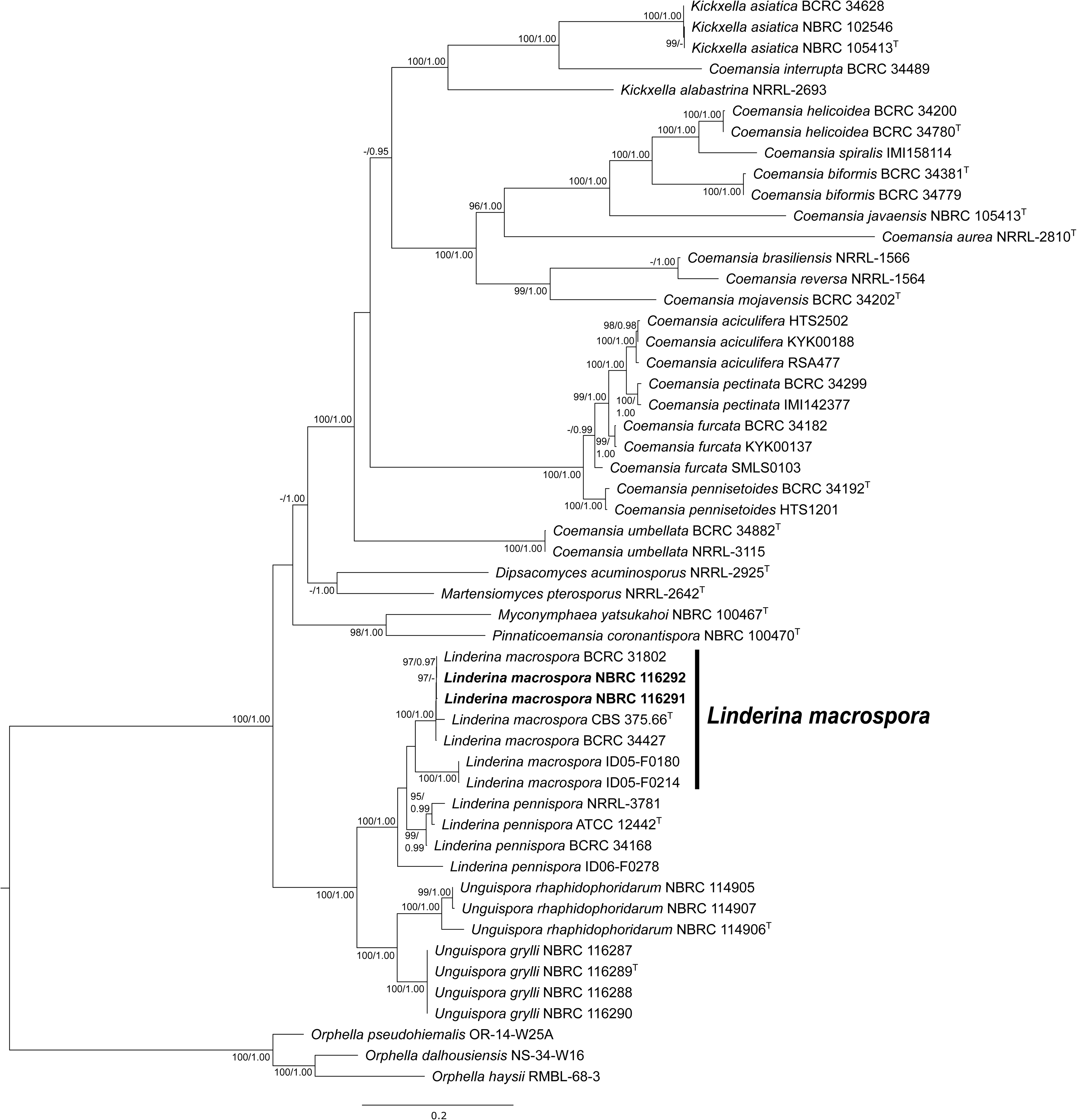
Maximum likelihood (ML) tree of *Kickxellales* based on SSU-ITS-LSU sequences. Number at the nodes indicate ML ultrafast bootstrap values (UFB) ≥ 95% and Bayesian posterior probabilities (PP) ≥ 0.95 as UFB/PP. Hyphens (-) represent values below thresholds or nodes not recovered in the respective analysis. Sequences obtained from our isolates are shown in bold. Strains derived from holotypes are indicated with a superscript T (^T^).

Based on the results of morphological observations and phylogenetic analyses, the examined isolates were identified as *L. macrospora*. A detailed morphological description is provided below:

*Linderina macrospora* Y. Chang (1967)

MycoBank No: MB333396

Figure. 1

Description: Colonies on 1/2 ME-YE agar pale yellow to yellow, nearly 1 cm high. Vegetative hyphae hyaline, septate, 2.5–10.5 μm wide. Sporangiophores yellowish, erect, septate, simple or occasionally branched irregularly, asperulate, 7–16(–18) μm wide, tapering above and terminating in a single sporocladium or producing occasionally two to three, rarely four to seven sporocladia sympodially on one sporangiophore. Sporocladia dome-shaped, sessile, aseptate, asperulate, (13–)13.5–22(–22.5) × (15.5–)16.5–25.5 μm, producing pseudophialides on the upper surface. Pseudophialides flask-shaped, asperulate, 8–14.5(–15.5) × 3–4 μm, bearing a sporangiolum apically. Sporangiola hyaline, cylindrical, tapered, swelling on the tip and truncate at the base, 26–30.5(–32.5) × 3.5–5 μm, containing a sporangiospore at the basal end, easily detached and immersed in a liquid droplet at maturity. Sporangiospores hyaline, lanceolate, uniformly tapered to the acute apices, round below, 22.5–27.5(–29.5) × 3.5–5 μm. Zygospores not observed.

Specimens examined: JAPAN, Ibaraki, Sakuragawa-shi, Tomiya, 36°22’44.3”N, 140°06’24.8”E, soil in the mixed forest of needleleaf and broadleaf trees, 29 Apr 2021, T. Ri, living culture NBRC 116291 (SSU = PZ418801, LSU = PZ425532, ITS = PZ418799); Ibaraki, Kuji-gun, Daigo, Konamase, 36°45’30”N, 140°27’39”E, soil in the woods, 20 Oct 2015, T. Masaki, living culture NBRC 116292 (SSU = PZ418802, LSU = PZ425533, ITS = PZ418800).

### 3.2. Influence of anaerobic conditions on the morphology of Linderina macrospora

Under anaerobic conditions on 1/2 ME-YE agar at 20 °C, the sporangiola formed arthrospores (Fig. 3A, B). The sporangiospores germinated at the basal end and swelled at the germination point beyond their original width, consistent with observations under aerobic conditions (Fig. 3C). However, the fungal thalli were distinguished from usual hyphae formed under aerobic conditions by being wider, more linear and branching at right angles (Fig. 3A, D). Segmentation occurred irregularly with cells connected or disarticulated from each other (Fig. 2E, F). Septa formed by the segmentation lacked a central pore with a plug (Fig. 3C, E, F). The size of the arthrospores varied widely due to the variability in the degree of hyphal elongation and frequency of segmentation (Fig. 3E, F). Additionally, we observed a gradation of morphogenesis, ranging from *Geotrichum*-like filamentous growth to yeast-like unicellular proliferation (Fig. 3A, B). The arthrospores germinated from the segmentation sites and multiplied by fission or budding.

**Figure 3.**
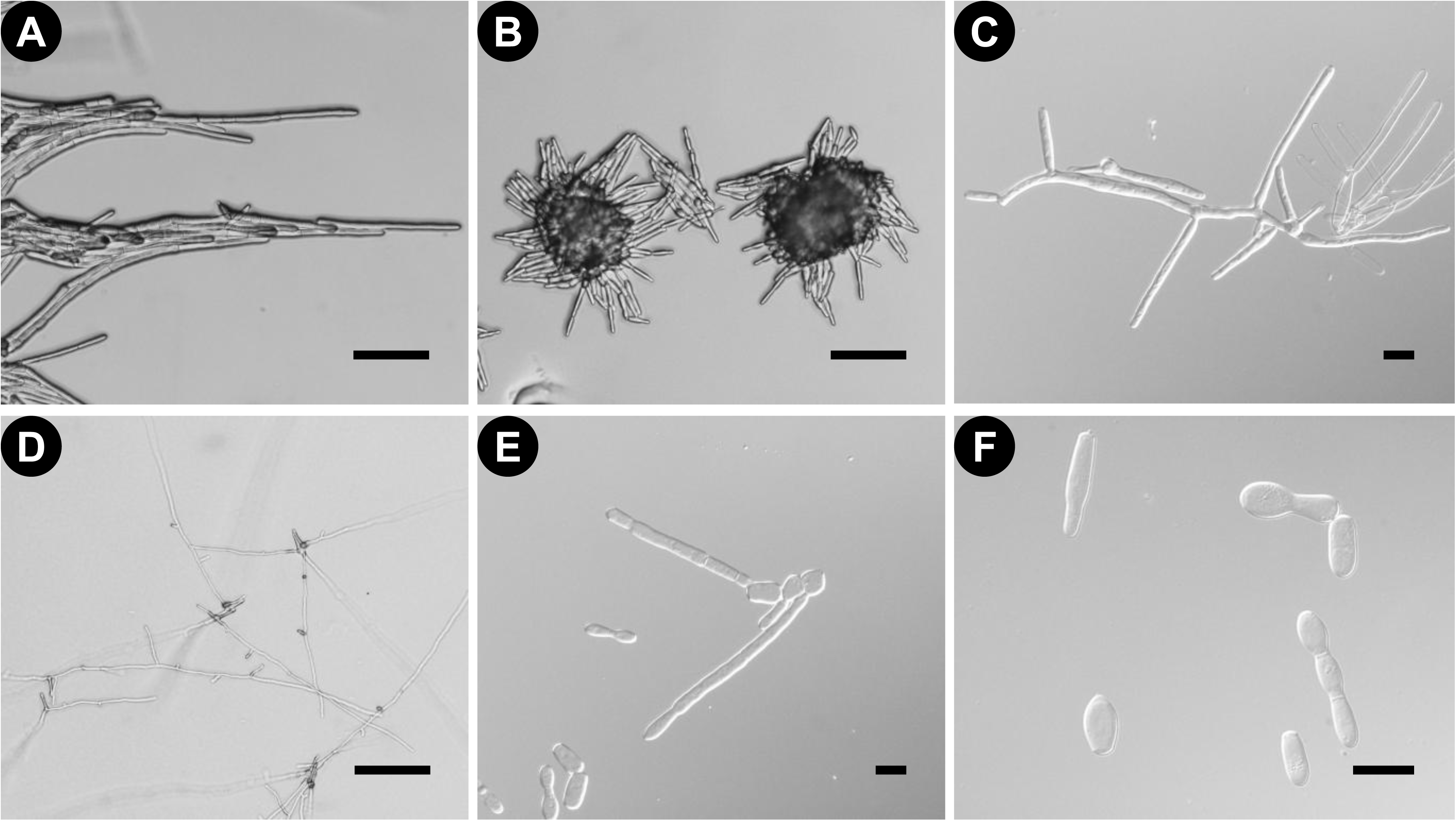
Anaerobic growth of *Linderina macrospora* (**A, C, E, F** NBRC 116291; **B, D** NBRC 116292). **C, E, F** are mounted in lactic acid. **A, E, F** are 7-day-old cultures incubated on 1/2 ME-YE agar at 20 °C. **B, C** are 2-day-old cultures incubated on 1/2 ME-YE agar at 20 °C. **D** is a 2-day-old culture incubated on CMA at 20 °C. **A** Arthrospore formation in the filamentous form like *Geotrichum*. **B** Arthrospore formation in the yeast-like form. **C** Germinated sporangiospores. **D** Hyphal growth without arthrospore formation. **E, F** Arthrospores. Scalebars: 100 μm (**A, B, D**); 20 μm (**C, E, F**).

The effects of medium and temperature on arthrospore formation are presented in Table 1. Arthrospore formation was observed on nearly all media tested, although the degree of arthrosporogenesis varied among the culture plates. In some cases, both arthrosporogenesis and formation of vegetative hyphae submerged in the media were observed concurrently (indicated as “w (weak)” in Table 1). In contrast, arthrospore formation on CMA was completely absent in NBRC 116291, which was the only instance among the medium conditions, whereas NBRC 116292 exhibited a mixture of arthrosporogenesis and normal hyphal growth. No distinct differences were observed between the 20 and 25 °C treatments. Under microaerophilic conditions, only normal hyphal growth occurred, regardless of the isolate.

**Table 1.** Effects of culture medium and temperature on anaerobic arthrospore formation in *Linderina macrospora* (three replicates).

| Isolate | Temperature | 1/2 ME-YE agar | -ME<sup>a</sup> | -YE<sup>a</sup> | -P<sup>a</sup> | -G<sup>a</sup> | PDA | CMA |
| --- | --- | --- | --- | --- | --- | --- | --- | --- |
| NBRC 116291 | 20 °C | + | +, w, - | +, w | + | +, w, - | + | - |
|  | 25 °C | +, w | +, w, - | +, w | + | + | +, w | - |
| NBRC 116292 | 20 °C | + | +, w | +, w | + | +, w | + | +, w, - |
|  | 25 °C | +, w | +, w | +, w | + | + | + | +, w, - |

+ = only arthrospore formation, w (weak) = both arthrospore formation and usual hyphal growth, - = only usual hyphal growth.
<sup>a</sup>Media by omitting individual components from 1/2 ME-YE agar: -ME (malt extract), -YE (yeast extract), -P (peptone), and -G (glucose).

### 3.3. Physiological characteristics of Linderina macrospora

The colony sizes of *L. macrospora* at each temperature are shown in Figure 4. Both isolates grew faster as the incubation temperature increased up to 30 °C. However, NBRC 116291 grew slower at 35 °C, comparable to that at 10 °C. Their optimum temperature for vegetative growth was between 25 °C and 30 °C. Sporulation was observed only at 20, 25, and 30 °C.

**Figure 4.**
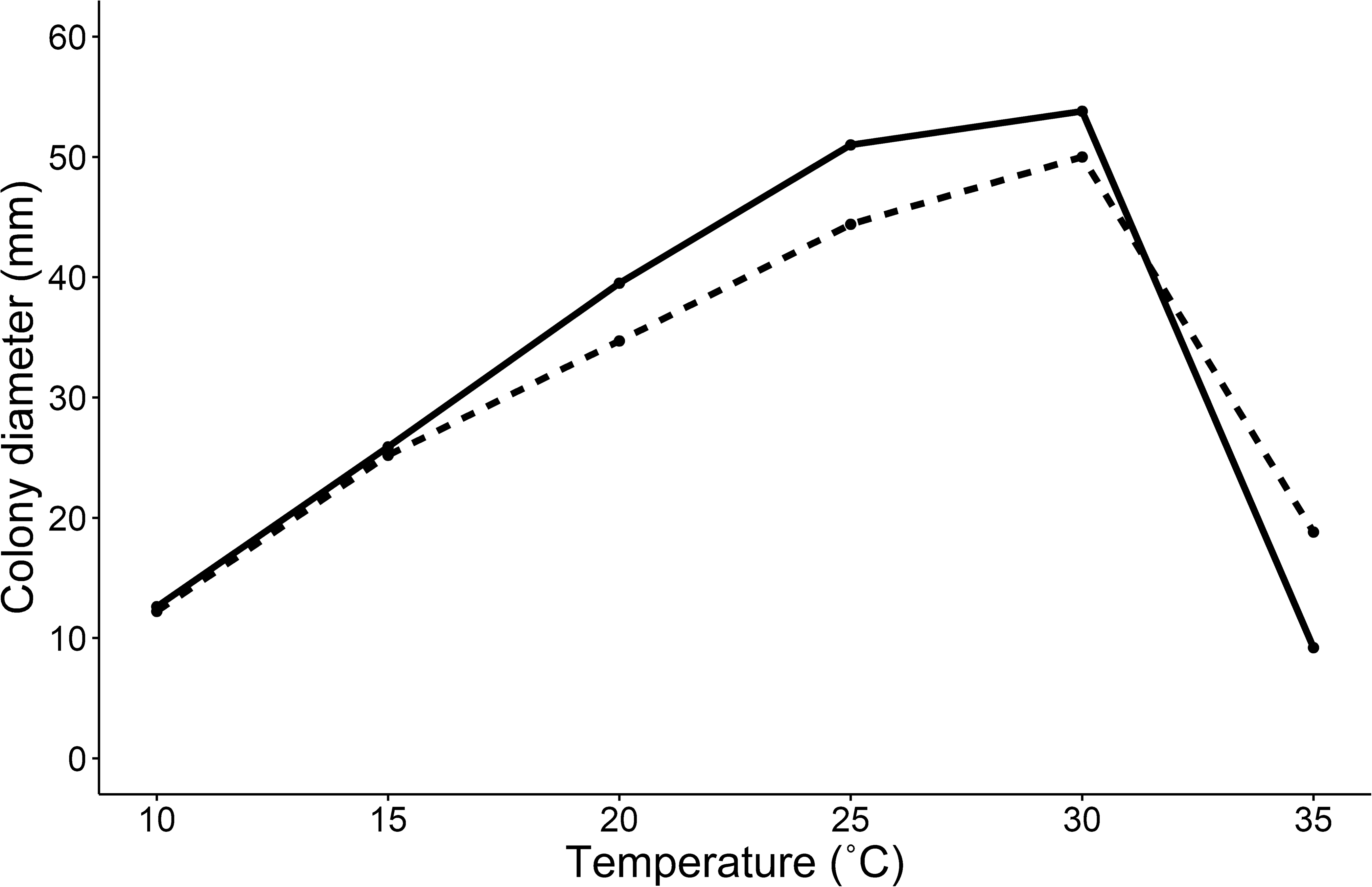
Mean colony diameter of *Linderina macrospora* cultured on CMA for 10 days at various temperatures. The solid and dashed lines represent NBRC116291 and NBRC 116292, respectively.

The effects of medium and illumination conditions on their growth and sporulation are summarized in Table 2. When comparing between nutrient-rich and nutrient-poor media under “dark” conditions, the growth and sporulation in nutrient-rich media were generally better for both isolates. Colonies were slightly larger and yellow in nutrient-rich media, whereas they appeared white in nutrient-poor media (Fig. 5A). Both vegetative hyphae and sporangiophores were more abundant, and spore production was visibly improved. However, the number of sporocladia per sporangiophore was greater in nutrient-poor media than in nutrient-rich media. It was also influenced by isolate and temperature and was greater especially when the conditions were NBRC 116292 and 25 °C, respectively. Compared to CMA and _LC_A, vegetative growth and sporangiophore production were better in _LC_A, and sporocladium production per sporangiophore was indistinguishable. Compared to PDA and 1/2 ME-YE agar, colony size was larger in PDA, but sporulation was unstable in PDA, such as at 20 °C.

**Figure 5.**
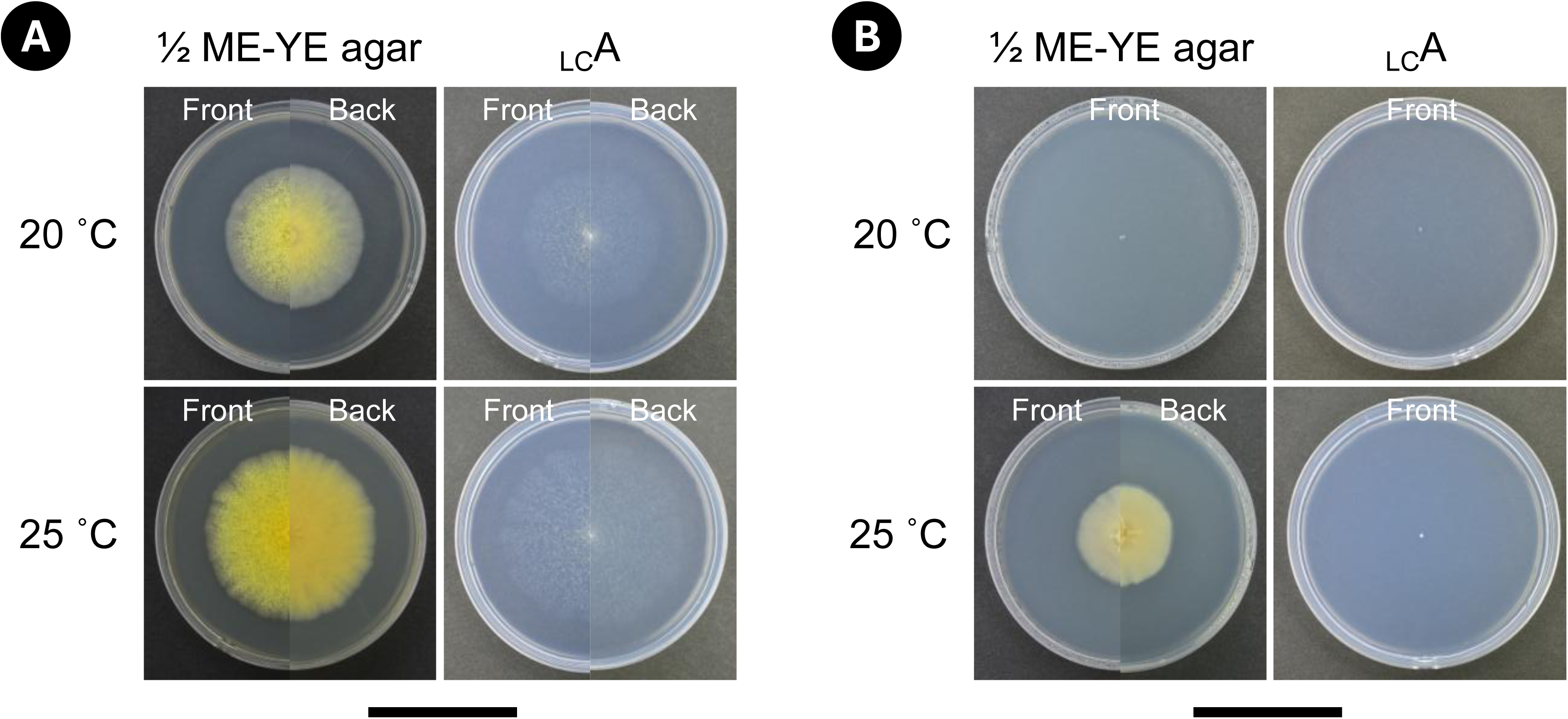
Comparison of colony appearance of *Linderina macrospora* (NBRC 116292) under different temperature, culture medium, and illumination conditions. **A** The 10-day cultures in constant darkness by shading with aluminum foil. **B** The 10-day cultures in constant light with white LED lamps. Scalebars: 5 cm (**A, B**).

**Table 2.**
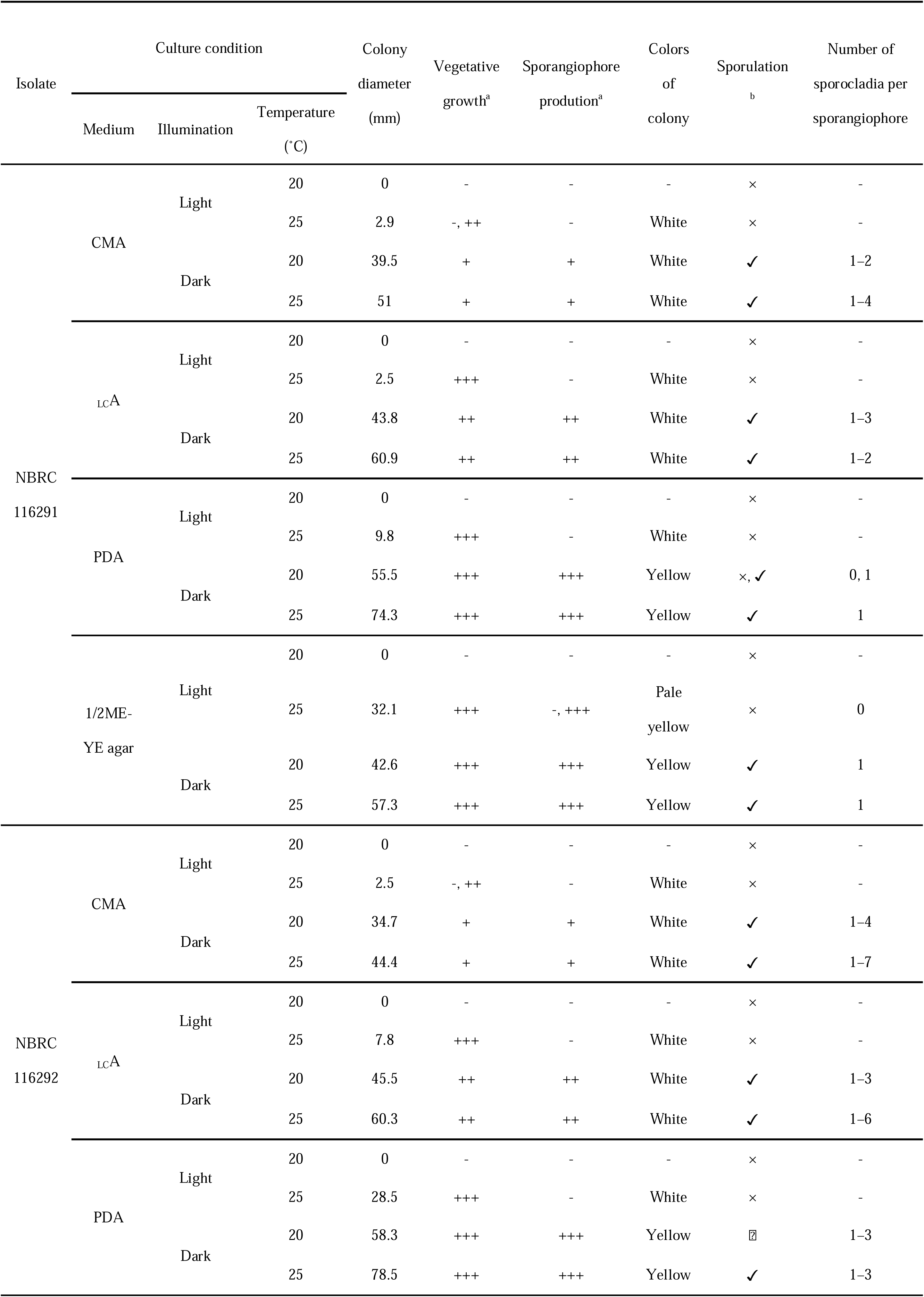
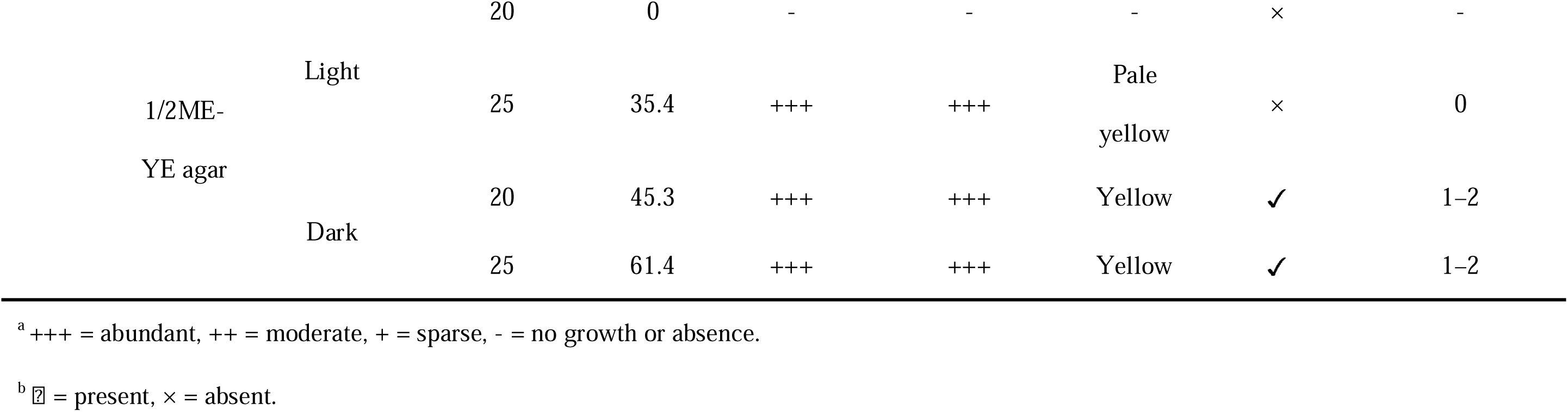
Influence of culture medium, illumination and temperature on the growth and asexual reproduction of two isolates of *Linderina macrospora*.

| Isolate | Culture condition |  |  | Colony diameter (mm) | Vegetative growth<sup>a</sup> | Sporangiophore production<sup>a</sup> | Colors of colony | Sporulation<sup>b</sup> | Number of sporocladia per sporangiophore |
| --- | --- | --- | --- | --- | --- | --- | --- | --- | --- |
|  | Medium | Illumination | Temperature (°C) |  |  |  |  |  |  |
| NBRC 116291 | CMA | Light | 20 | 0 | - | - | - | × | - |
|  |  |  | 25 | 2.9 | -, ++ | - | White | × | - |
|  |  | Dark | 20 | 39.5 | + | + | White | ✓ | 1–2 |
|  |  |  | 25 | 51 | + | + | White | ✓ | 1–4 |
|  | LCA | Light | 20 | 0 | - | - | - | × | - |
|  |  |  | 25 | 2.5 | +++ | - | White | × | - |
|  |  | Dark | 20 | 43.8 | ++ | ++ | White | ✓ | 1–3 |
|  |  |  | 25 | 60.9 | ++ | ++ | White | ✓ | 1–2 |
|  | PDA | Light | 20 | 0 | - | - | - | × | - |
|  |  |  | 25 | 9.8 | +++ | - | White | × | - |
|  |  | Dark | 20 | 55.5 | +++ | +++ | Yellow | ×, ✓ | 0, 1 |
|  |  |  | 25 | 74.3 | +++ | +++ | Yellow | ✓ | 1 |
|  | 1/2ME-YE agar | Light | 20 | 0 | - | - | - | × | - |
|  |  |  | 25 | 32.1 | +++ | -, +++ | Pale yellow | × | 0 |
|  |  | Dark | 20 | 42.6 | +++ | +++ | Yellow | ✓ | 1 |
|  |  |  | 25 | 57.3 | +++ | +++ | Yellow | ✓ | 1 |
| NBRC 116292 | CMA | Light | 20 | 0 | - | - | - | × | - |
|  |  |  | 25 | 2.5 | -, ++ | - | White | × | - |
|  |  | Dark | 20 | 34.7 | + | + | White | ✓ | 1–4 |
|  |  |  | 25 | 44.4 | + | + | White | ✓ | 1–7 |
|  | LCA | Light | 20 | 0 | - | - | - | × | - |
|  |  |  | 25 | 7.8 | +++ | - | White | × | - |
|  |  | Dark | 20 | 45.5 | ++ | ++ | White | ✓ | 1–3 |
|  |  |  | 25 | 60.3 | ++ | ++ | White | ✓ | 1–6 |
|  | PDA | Light | 20 | 0 | - | - | - | × | - |
|  |  |  | 25 | 28.5 | +++ | - | White | × | - |
|  |  | Dark | 20 | 58.3 | +++ | +++ | Yellow | ☒ | 1–3 |
|  |  |  | 25 | 78.5 | +++ | +++ | Yellow | ✓ | 1–3 |

|  |  |  |  |  |  |  |  |  |
| --- | --- | --- | --- | --- | --- | --- | --- | --- |
| 1/2ME-YE agar | Light | 20 | 0 | - | - | - | × | - |
|  |  | 25 | 35.4 | +++ | +++ | Pale<br>yellow | × | 0 |
|  | Dark | 20 | 45.3 | +++ | +++ | Yellow | ✓ | 1-2 |
|  |  | 25 | 61.4 | +++ | +++ | Yellow | ✓ | 1-2 |

<sup>a</sup> +++ = abundant, ++ = moderate, + = sparse, - = no growth or absence.
<sup>b</sup> ☐ = present, × = absent.

Under “light” conditions, mycelial growth and sporulation of *L. macrospora* were inhibited in all the treatments. At 20 °C, sporangiospores partially germinated, but the isolates eventually lost viability without further growth (Fig. 5B). At 25 °C, they grew more or less but failed to sporulate (Fig. 5B). The colony diameters after 10 days of incubation were less than 10 mm in nutrient-poor media against around 30 mm in nutrient-rich media. NBRC 116292 grew better than the other isolate. The growth inhibition was also observed under a 14:10 h light/dark cycle with white light. The range of colony diameters after 10 days was 0–10 mm at 20 °C and 22–32 mm at 25 °C. The colonies incubated at 25 °C showed a concentric pattern of growth. In a replicate on CMA at 20 °C, spore germination was arrested, resulting in a loss of viability. Sporulation was almost entirely absent; although it was observed on a single plate on CMA at 25 °C, only two sporocladia and their sporangiola were produced (the viability of these sporangiola was subsequently confirmed). In these experiments, submerged mycelia and sporangiophores exhibited no phototropic responses.

## 4. Discussion

Arthrospore formation was newly identified in *Linderina* in the present study, whereas it is already known to occur in *Pinnaticoemansia coronantispora* among kickxellalean fungi (Kurihara and Degawa, 2006). However, these genera differ in their morphogenetic patterns and the factors that trigger arthrospore formation. In *P. coronantispora*, sporangiospores germinate downward and undergo repeated dichotomy under aerobic, nutrient-rich conditions (Kurihara and Degawa, 2006). In contrast, arthrospores of *L. macrospora* are produced through irregular segmentation under anaerobic, nutrient-rich conditions (Fig. 3A–C, E, F). The degree of thallic elongation and the size of arthrospores also vary more widely in *L. macrospora* than in *P. coronantispora*. During arthrospore formation, the thallus morphology of *L. macrospora* exhibited a continuum from filamentous to yeast-like forms (Fig. 3A, B), which appeared to be highly sensitive to subtle environmental gradients within the same culture conditions. In *Unguispora*, sporangiospores germinate basally and elongate downward to form secondary spores via septation, as in these genera; however, the secondary spores subsequently swell while remaining attached to the sporangiospores (Ri et al., 2022). The formation of arthrospores in *Pinnaticoemansia* and secondary spores in *Unguispora* is regarded as a form of yeast-like growth, characterized by single-cell multiplication and determinate growth. Although arthrospore formation in *L. macrospora* appears to be environmentally labile, unicellular proliferation via fission or budding was observed (Fig. 3B, F), which can also be classified as yeast-like growth. Unlike these genera, *Coemansia*, a common saprobic fungus, has not been found to grow in any form other than hyphae, according to previous studies (Benny et al., 2016) and our observations. Therefore, the occurrence of arthrospore formation in *L. macrospora* suggests its potential ecological relationship with the animal gut in nature.

Atmospheric conditions appeared to be a critical trigger for arthrospore formation in *L. macrospora*. Our results showed that arthrospore formation occurred under anaerobic conditions (less than 0.1% O_2_ and more than 15% CO_2_) but was absent under microaerophilic conditions (6–12% O_2_ and 5–8% CO_2_). This highlights the need to distinguish whether low oxygen or elevated carbon dioxide is the primary driver and to determine their respective concentration thresholds. Furthermore, the responsiveness to the environmental triggers differed between the isolates. For instance, NBRC 116292 exhibited a higher propensity for arthrospore formation on CMA than NBRC 116291 (Table 1), suggesting that individual isolates possess different sensitivity thresholds for nutritional stimuli. Regarding the nutritional basis for this response, the metabolic switch to arthrospore formation may be governed by the availability of nitrogen sources or trace elements rather than carbon concentration. This is supported by our observation that, while nutrient-poor media were generally unsuitable for the process, the specific omission of glucose did not hinder it (Table 1). Notably, contrary to our observations, the only previous study on anaerobic cultivation did not consider *Linderina* as an anaerobic fungus (Hesseltine et al., 1985). That study utilized *L. pennispora* instead of *L. macrospora*, which was used in this study. Although slight mycelial growth was observed after seven days of anaerobic incubation on PDA at 25°C using GasPak, *L. pennispora* was ultimately classified as an obligate aerobe based on growth tests under different atmospheric conditions using CDC anaerobe blood agar (Hesseltine et al., 1985). Combining the previous report with our findings raise two alternative possibilities regarding the reported absence of arthrospores: either the capacity for arthrospore formation is species-specific within *Linderina*, or the previous investigation overlooked this development owing to methodological limitations. Supporting the latter, our observations confirmed that arthrospore formation and vegetative mycelial growth occurred concurrently in *L. macrospora* (Table 1). Thus, previous investigations using stereomicroscopy might have missed arthrospores or distinctive thalli. In view of the variation in responsiveness between isolates, arthrospore formation in *Linderina* warrants re-evaluation using a broader, multi-species sampling.

This study integrates previously fragmented reports on the cultural characteristics of *L. macrospora*. The temperature test in this study demonstrated that the optimal temperature for both vegetative growth and sporulation lies between 25 and 30 °C (Fig. 4). This finding is consistent with previous reports that utilized incubation temperatures of 25 and 26°C (Chang, 1967; Kurihara et al., 2008). Regarding the comparison of culture media, the nutrient-rich media were superior in terms of vegetative growth and total spore yield (Table 2). However, the nutrient-poor media resulted in a higher number of sporocladia per sporangiophore. In NBRC116291, the number of sporocladia per sporangiophore was two when observed in the incubated soil, whereas it decreased to one on nutrient-rich media (Table 2). This is likely because resources were preferentially allocated to rapid vegetative growth, resulting in the morphological simplification of reproductive structures. This trend also holds true for *L. pennispora*, as evidenced by the culture results on CMA and PDA (Ho et al., 2007). Although some previous studies have noted that nutrient-poor conditions favor sporulation in *Linderina* (Chuang and Ho, 2009; Kurihara et al., 2008), they did not explicitly distinguish between the total spore count and the density of sporocladia on a sporangiophore, and it appears that prior claims may have been based on the latter. As the number of sporocladia per sporangiophore is a critical diagnostic characteristic for distinguishing species within *Linderina*, our results highlight the necessity of conducting morphological evaluations on nutrient-poor media to ensure taxonomic accuracy.

In addition, exposure to white light had a profound impact on the growth and development of *L. macrospora* (Fig. 5, Table 2). The observed effects varied with temperature: at 20 °C, vegetative growth was severely inhibited, often leading to cell death, whereas at 25 °C, vegetative growth occurred but was consistently accompanied by a failure in sporulation. Both continuous light and long-day conditions (14-hour photoperiod) resulted in diminished vegetative growth and sporulation deficiency, suggesting that this species is poorly adapted to environments with light exposure. To date, Kurihara et al. (2008) are the only researchers who have explicitly specified light conditions, utilizing a 15:9 light-dark cycle. They reported that *L. macrospora* exhibited sparse sporulation in media lacking invertebrate-derived substrates, requiring even several months for induction. Other studies have not documented low sporulation or a requirement for months-long incubation for sporulation in *L. macrospora*. Therefore, these earlier findings may also reflect the light-induced inhibition of sporulation, as observed in the present study. Considering that their Indonesian isolates are phylogenetically distinct from our isolates (Fig. 2), it is plausible that this light-responsiveness is a more broadly shared trait rather than an exceptional case limited to specific strains. Furthermore, although we have observed positive phototropism in the sporangiophores of *Unguispora* (unpublished data), light-dependent developmental regulation has not been reported in the other genera within *Kickxellales*. It remains to be determined whether this extreme light sensitivity is a unique trait of *L. macrospora* or *Linderina*, highlighting the need for broader comparative studies across *Kickxellales*.

Many fungi respond to visible light, which regulates diverse processes, such as sporulation, germination, and secondary metabolism (Fuller et al., 2015; Yu and Fischer, 2019). While light-induced inhibition of sporulation has been reported, our findings revealed an exceptionally hypersensitive response in the species. For instance, while white LEDs at a photon flux density comparable to ours completely inhibit sporulation in many isolates of *Botrytis cinerea*, this inhibitory effect is not universal across all tested isolates (Meng et al., 2020). In *Aspergillus fumigatus*, it inhibits conidial germination but has little effect on conidial production (Fuller et al., 2013). Furthermore, white light restricts the growth of *A. ochraceus* and *A. oryzae* without stopping it completely (Hatakeyama et al., 2007; Murthy et al., 2015; Zhang et al., 2021). In contrast, *L*. macrospora is far more severe and comprehensive; it exhibited a virtually complete loss of sporulation, coupled with an unprecedented lethal growth inhibition at 20 °C. The consistent suppression of both sporulation and vegetative growth under low-intensity light underscores a physiologically unique trait. These results suggest that the ecological niche of *L. macrospora* is effectively restricted to light-shielded habitats. Accordingly, the species may rely primarily on passive transport within the soil as its principal mode of dispersal. Moreover, our observations of arthrospore formation, indicative of adaptation to low oxygen conditions, further suggest dispersal via ingestion by soil fauna. Given the lack of research on photoresponses in *Kickxellomycotina* (Corrochano and Garre, 2010; Idnurm et al., 2010), elucidating the molecular mechanisms of this photophysiology will provide critical insights into the evolutionary adaptation of fungi to light.

## 5. Conclusions

This study represents the first report of *L. macrospora* from Japan. Our culture examination revealed that the optimal growth temperature for this species lies between 25°C and 30°C, providing essential data for both culture optimization and potential distribution modeling. Furthermore, it demonstrated that nutrient-poor media, such as CMA, are most suitable for morphological observation, establishing a standardized methodological approach for taxonomy. Notably, this research uncovered two distinct physiological traits of *L. macrospora*: arthrospore formation under anaerobic, nutrient-rich conditions and the suppression of both vegetative growth and sporulation by light. The formation of arthrospores, which occasionally leads to a yeast-like colony appearance, suggests a possible ecological link to animal intestinal tracts, similar to the genus *Unguispora*. Moreover, exposure to white light resulted in the complete inhibition of sporulation across nearly all treatments; notably, at 20°C, light exposure was lethal. These findings strongly suggest that the natural habitat of this fungus is restricted to light-shielded environments, such as deep soil layers. The discovery of these specialized physiological characteristics underscores the necessity of culture-based studies in overcoming current phylogenetic bottlenecks. This approach warrants further screening for similar traits across the *Kickxellales*, which will be essential for a holistic understanding of their evolutionary biology.

## CRediT authorship contribution statement

**Tomohiko Ri**: Conceptualization, Investigation, Writing – Original Draft, Writing – Review & Editing, Data Curation, Funding Acquisition, Resources. **Teruhisa Masaki**: Resources, Writing – Review & Editing. **Yousuke Degawa**: Resources, Supervision, Writing – review & editing.

## Declarations

The authors declare that there is no ethical conflict. This article does not contain any studies with animals performed by any of the authors.

## Acknowledgments

This study was supported, in part, by a Grant-in-Aid for Scientific Research (no. 24KJ0485, 26KJ0180 to T. R.) from the Japan Society for the Promotion of Sciences. We thank Masana Izawa for cooperating to collect soil samples.

## References

Baijal, U., 1963. *Linderina pennispora* Raper & Fennell from India. Mycopathologia et Mycologia Applicata 21, 109–111. 10.1007/BF02049171

Benjamin, R.K., 1961. Addenda to “The merosporangiferous *Mucorales*.” aliso 5, 11–19. 10.5642/aliso.19610501.05

Benjamin, R.K., 1959. The merosporangiferous *Mucorales*. Aliso 4, 321–433. 10.5642/aliso.19590402.05

Benny, G.L., Humber, R.A., Morton, J.B., 2001. *Zygomycota*: *Zygomycetes*, in: McLaughlin, D.J., McLaughlin, E.G., Lemke, P.A. (Eds.), The Mycota, Vol 7. Systematics and Evolution Part A. Springer-Verlag, Berlin, pp. 113–146. 10.1007/978-3-662-10376-0

Benny, G.L., Humber, R.A., Voigt, K., 2014. 8 Zygomycetous fungi: phylum *Entomophthoromycota* and subphyla *Kickxellomycotina*, *Mortierellomycotina*, *Mucoromycotina*, and *Zoopagomycotina*, in: McLaughlin, D.J., Spatafora, J.W. (Eds.), Systematics and Evolution: Part A, The Mycota. Springer, Berlin, Heidelberg, pp. 209–250. 10.1007/978-3-642-55318-9_8

Benny, G.L., Smith, M.E., Kirk, P.M., Tretter, E.D., White, M.M., 2016. Challenges and future perspectives in the systematics of *Kickxellomycotina*, *Mortierellomycotina*, *Mucoromycotina*, and *Zoopagomycotina*, in: Li, D.-W. (Ed.), Biology of Microfungi, Fungal Biology. Springer International Publishing, Switzerland, pp. 65–126. 10.1007/978-3-319-29137-6_5

Capella-Gutiérrez, S., Silla-Martínez, J.M., Gabaldón, T., 2009. trimAl: a tool for automated alignment trimming in large-scale phylogenetic analyses. Bioinformatics 25, 1972–1973. 10.1093/bioinformatics/btp348

Carmichael, J.W., 1955. Lacto-fuchsin: a new medium for mounting fungi. Mycologia 47, 611. 10.1080/00275514.1955.12024478

Chang, Y., 1967. *Linderina macrospora* sp. nov. from Hong Kong. Transactions of the British Mycological Society 50, 311–314. 10.1016/S0007-1536(67)80043-3

Chernomor, O., von Haeseler, A., Minh, B.Q., 2016. Terrace aware data structure for phylogenomic inference from supermatrices. Systematic Biology 65, 997–1008. 10.1093/sysbio/syw037

Chien, C.-Y., 1971. *Linderina macrospora* from forest soil of the southeastern United States. Mycologia 63, 410–412. 10.1080/00275514.1971.12019119

Chuang, S.-C., Ho, H.-M., 2009. Notes on *Zygomycetes* of Taiwan (VII): two kickxellalean species, Linderina macrospora and Ramicandelaber brevisporus new to Taiwan. Fungal Science 24, 23–28. 10.7099/FS.200912.0023

Chuang, S.-C., Ho, H.-M., Reynolds, N., Smith, M.E., Benny, G.L., Chien, C.-Y., Tsai, J.-L., 2017. Preliminary phylogeny of *Coemansia* (*Kickxellales*), with descriptions of four new species from Taiwan. Mycologia 109, 815–831. 10.1080/00275514.2017.1401892

Corrochano, L.M., Garre, V., 2010. Photobiology in the *Zygomycota*: multiple photoreceptor genes for complex responses to light. Fungal Genetics and Biology, Special Issue: Photobiology 47, 893–899. 10.1016/j.fgb.2010.04.007

Doweld, A., 2014a. Nomenclatural novelties: Ramicandelaberaceae fam. nov. and Ramicandelaberales ord. nov. Index Fungorum.

Doweld, A., 2014b. Nomenclatural novelties: Spiromycetaceae fam. nov. and Spiromycetales ord. nov. Index Fungorum.

Fuller, K.K., Loros, J.J., Dunlap, J.C., 2015. Fungal photobiology: visible light as a signal for stress, space and time. Curr Genet 61, 275–288. 10.1007/s00294-014-0451-0

Fuller, K.K., Ringelberg, C.S., Loros, J.J., Dunlap, J.C., 2013. The fungal pathogen *Aspergillus fumigatus* regulates growth, metabolism, and stress resistance in response to light. mBio 4, 10.1128/mbio.00142-13. 10.1128/mbio.00142-13

Gardes, M., Bruns, T.D., 1993. ITS primers with enhanced specificity for basidiomycetes - application to the identification of mycorrhizae and rusts. Molecular Ecology 2, 113–118. 10.1111/j.1365-294X.1993.tb00005.x

Hatakeyama, R., Nakahama, T., Higuchi, Y., Kitamoto, K., 2007. Light represses conidiation in koji mold *Aspergillus oryzae*. Bioscience, Biotechnology, and Biochemistry 71, 1844–1849. 10.1271/bbb.60713

Hesseltine, C.W., Featherston, C.L., Lombard, G.L., Dowell, V.R., 1985. Anaerobic growth of molds isolated from fermentation starters used for foods in Asian countries. Mycologia 77, 390–400. 10.2307/3793195

Ho, H.-M., Chien, C.-Y., Chuang, S.-C., 2007. Notes on *Zygomycetes* of Taiwan (V): *Linderina pennispora* new to Taiwan. Fungal Science 22, 35–38.

Hoang, D.T., Chernomor, O., von Haeseler, A., Minh, B.Q., Vinh, L.S., 2018. UFBoot2: Improving the ultrafast bootstrap approximation. Molecular Biology and Evolution 35, 518–522. 10.1093/molbev/msx281

Idnurm, A., Verma, S., Corrochano, L.M., 2010. A glimpse into the basis of vision in the kingdom *Mycota*. Fungal Genet Biol 47, 881–892. 10.1016/j.fgb.2010.04.009

Jackson, H.S., Dearden, E.R., 1948. *Martensella corticii* Thaxter and its distribution. Mycologia 40, 168–176. 10.1080/00275514.1948.12017697

Kalyaanamoorthy, S., Minh, B.Q., Wong, T.K.F., von Haeseler, A., Jermiin, L.S., 2017. ModelFinder: fast model selection for accurate phylogenetic estimates. Nature Methods 14, 587–589. 10.1038/nmeth.4285

Katoh, K., Rozewicki, J., Yamada, K.D., 2019. MAFFT online service: multiple sequence alignment, interactive sequence choice and visualization. Briefings in Bioinformatics 20, 1160–1166. 10.1093/bib/bbx108

Kurihara, Y., Degawa, Y., 2006. *Pinnaticoemansia*, a new genus of *Kickxellales*, with a revised key to the genera of *Kickxellales*. Mycoscience 47, 205–211. 10.1007/S10267-006-0294-8

Kurihara, Y., Park, J.-Y., Ando, K., Sukarno, N., Ilyas, M., Yuniarti, E., Saraswati, R., Mangunwardoyo, W., Widyastuti, Y., 2008. Indonesian *Kickxellales*: two species of *Coemansia* and *Linderina*. Mycoscience 49, 250–257. 10.1007/S10267-008-0417-5

Kurihara, Y., Tokumasu, S., Chien, C.-Y., 2000. *Coemansia furcata* sp. nov. and its distribution in Japan and Taiwan. Mycoscience 41, 579–583. 10.1007/BF02460924

Loh, L.S., Nawawi, A., Kuthubutheen, A.J., 2001. Mucoraceous fungi from Malaysia. Institute of Biological Sciences, University of Malaya, Kuala Lumpur.

Meng, L., Mestdagh, H., Ameye, M., Audenaert, K., Höfte, M., Van Labeke, M.-C., 2020. Phenotypic variation of *Botrytis cinerea* isolates is influenced by spectral light quality. Front. Plant Sci. 11. 10.3389/fpls.2020.01233

Miura, K., Kudo, M., 1970. An agar-medium for aquatic hyphomycetes [In Japanese]. Trans. Mycol. Soc. Japan 11, 116–118.

Murthy, P.S., Suzuki, S., Kusumoto, K.-I., 2015. Effect of light on the growth and acid protease production of *Aspergillus oryzae*. Food Science and Technology Research 21, 631–635. 10.3136/fstr.21.631

O’Donnell, K., Cigelnik, E., Benny, G.L., 1998. Phylogenetic Relationships among the *Harpellales* and *Kickxellales*. Mycologia 90, 624–639. 10.2307/3761222

Rambaut, A., Drummond, A.J., Xie, D., Baele, G., Suchard, M.A., 2018. Posterior summarization in Bayesian phylogenetics using Tracer 1.7. Syst Biol 67, 901–904. 10.1093/sysbio/syy032

Raper, K.B., Fennell, D.I., 1952. Two noteworthy fungi from Liberian soil. American Journal of Botany 39, 79–86. 10.2307/2438097

Reynolds, N.K., Stajich, J.E., Benny, G.L., Barry, K., Mondo, S., LaButti, K., Lipzen, A., Daum, C., Grigoriev, I.V., Ho, H.-M., Crous, P.W., Spatafora, J.W., Smith, M.E., 2023. Mycoparasites, gut dwellers, and saprotrophs: phylogenomic reconstructions and comparative analyses of *Kickxellomycotina* fungi. Genome Biology and Evolution 15, evac185. 10.1093/gbe/evac185

Ri, T., Degawa, Y., 2025. *Unguispora grylli*, a new species of amphibious fungi associated with crickets (*Gryllidae*), transforms attachment structures of sporangiola in the host gut. Mycoscience 66, 162–170. 10.47371/mycosci.2025.01.001

Ri, T., Suyama, M., Takashima, Y., Seto, K., Degawa, Y., 2022. A new genus *Unguispora* in *Kickxellales* shows an intermediate lifestyle between saprobic and gut-inhabiting fungi. Mycologia 114, 934–946. 10.1080/00275514.2022.2111052

Ronquist, F., Teslenko, M., van der Mark, P., Ayres, D.L., Darling, A., Höhna, S., Larget, B., Liu, L., Suchard, M.A., Huelsenbeck, J.P., 2012. MrBayes 3.2: Efficient Bayesian phylogenetic inference and model choice across a large model space. Syst Biol 61, 539–542. 10.1093/sysbio/sys029

Schwarz, G., 1978. Estimating the dimension of a model. The Annals of Statistics 6, 461–464. 10.1214/aos/1176344136

Sugiura, N., 1978. Further analysis of the data by Akaike’s information criterion and the finite corrections: Further analysis of the data by akaike’ s. Communications in Statistics - Theory and Methods 7, 13–26. 10.1080/03610927808827599

Takashima, Y., Suyama, M., Yamamoto, K., Ri, T., Narisawa, K., Degawa, Y., 2022. Revisiting the isolation source after the first discovery: *Myconymphaea yatsukahoi* on excrements of *Lithobiomorpha* (*Chilopoda*). Mycoscience 63, 176–180. 10.47371/mycosci.2022.04.003

Tanabe, A.S., 2011. Kakusan4 and Aminosan: two programs for comparing nonpartitioned, proportional and separate models for combined molecular phylogenetic analyses of multilocus sequence data. Molecular Ecology Resources 11, 914–921. 10.1111/j.1755-0998.2011.03021.x

Tretter, E.D., Johnson, E.M., Benny, G.L., Lichtwardt, R.W., Wang, Y., Kandel, P., Novak, S.J., Smith, J.F., White, M.M., 2014. An eight-gene molecular phylogeny of the *Kickxellomycotina*, including the first phylogenetic placement of *Asellariales*. Mycologia 106, 912–935. 10.3852/13-253

Vilgalys, R., Hester, M., 1990. Rapid genetic identification and mapping of enzymatically amplified ribosomal DNA from several *Cryptococcus* species. Journal of Bacteriology 172, 4238–4246. 10.1128/jb.172.8.4238-4246.1990

Wong, T.K.F., Ly-Trong, N., Ren, H., Demotte, P., Baños, H., Roger, A.J., Susko, E., Bielow, C., De Maio, N., Goldman, N., Hahn, M.W., dos Reis, M., Vinh, L.S., Huttley, G., Lanfear, R., Minh, B.Q., 2026. IQ-TREE 3: phylogenomic inference software using complex evolutionary models. Mol Biol Evol 43, msag117. 10.1093/molbev/msag117

Yu, Z., Fischer, R., 2019. Light sensing and responses in fungi. Nat Rev Microbiol 17, 25–36. 10.1038/s41579-018-0109-x

Zhang, H., Wang, G., Yang, Q., Yang, X., Zheng, Y., Liu, Y., Xing, F., 2021. Effects of light on the ochratoxigenic fungi *Aspergillus ochraceus* and *A. carbonarius*. Toxins 13, 251. 10.3390/toxins13040251

